# Novel metrics for quantifying bacterial genome composition skews

**DOI:** 10.1101/176370

**Authors:** Lena M. Joesch-Cohen, Max Robinson, Neda Jabbari, Christopher Lausted, Gustavo Glusman

## Abstract

**Background:** Bacterial genomes have characteristic compositional skews, which are differences in nucleotide frequency between the leading and lagging DNA strands across a segment of a genome. It is thought that these strand asymmetries arise as a result of mutational biases and selective constraints, particularly for energy efficiency. Analysis of compositional skews in a diverse set of bacteria provides a comparative context in which mutational and selective environmental constraints can be studied. These analyses typically require finished and well-annotated genomic sequences.

**Results:** We present three novel metrics for examining genome composition skews; all three metrics can be computed for unfinished or partially-annotated genomes. The first two metrics, (dot-skew and cross-skew) depend on sequence and gene annotation of a single genome, while the third metric (residual skew) highlights unusual genomes by subtracting a GC content-based model of a library of genome sequences. We applied these metrics to all 7738 available bacterial genomes, including partial drafts, and identified outlier species. A number of these outliers (i.e., Borrelia, Ehrlichia, Kinetoplastibacterium, and Phytoplasma) display similar skew patterns despite only distant phylogenetic relationship. While unrelated, some of the outlier bacterial species share lifestyle characteristics, in particular intracellularity and biosynthetic dependence on their hosts.

**Conclusions:** Our novel metrics appear to reflect the effects of biosynthetic constraints and adaptations to life within one or more hosts on genome composition. We provide results for each analyzed genome, software and interactive visualizations at http://db.systemsbiology.net/gestalt/skew_metrics.

## Background

Bacterial genomes display significant compositional biases, both in terms of G+C content and in compositional skews, i.e., strand asymmetries in ‘T’ vs. ‘A’ and ‘G’ vs. ‘C’ usage [1]. These biases are proposed to arise from the complex interplay of differential mutation rates and multiple selective constraints [2, 3], particularly for energy efficiency [4], involving the replication, repair, and transcription enzymes. Bacterial chromosomes are replicated in both directions, from the origin of replication site to the terminator site; the “leading” strand is replicated continuously while the “lagging” strand is replicated in segments by different enzymes. Some genes are transcribed in the same direction as they are replicated (“leading strand genes”) while others are transcribed in the reverse direction (“lagging strand genes”). Each enzyme mediates both mutational and selective constraints, resulting in different compositional biases in different replicative and translational contexts. Analyses of skews in each context have the potential to expose multiple compositional constraints and their interactions, and ultimately inform about the DNA repair capacity, metabolism, and lifestyle of the species [5].

Compositional bias and strand asymmetry have been measured and analyzed in a variety of ways and contexts (for a review, see e.g. [5]). These methods include the original definitions (GC skew, (G-C)/(G+C); AT skew, (A-T)/(A+T)) [6], slight variants (e.g. G/(G+C)) [1], variants based on the three independent axes of Z Curves (x=R-Y, y=M-K, and z=S-W) [7, 8], ANOVA [9], correspondence analysis of codon bias metrics [10, 11], and competing mutational and selective parameters in an explicit evolutionary model [4], and have involved comparison of leading versus lagging contexts, transcribed versus intergenic regions, and restriction to each codon position.

Early examples of extreme compositional biases and asymmetries were found among species in the family Borreliaceae, tick-borne spirochetes including species causing Lyme disease (genus *Borreliella*, formerly *Borrelia*) as well as relapsing fever (genus *Borrelia*) [12]. Since its discovery in 1982 [13], the *Borreliella burgdorferi* spirochete has been of particular interest in the United States as the primary causative agent of Lyme disease. The sequencing of *B. burgdorferi* B31 in 1997 allowed an in depth exploration of the many intriguing features of the genome of this bacterium, from its small size and unusual structure (one large linear chromosome, several linear and circular plasmids) to its very low G+C content [14].

Significant skews in the third position of codons have been reported on both the leading (increased G and T) and lagging (increased A and C) strands in *B. burgdorferi* [10, 15]. Among the first 43 genomes investigated [1], *B. burgdorferi* had the most extreme difference between leading and lagging strand nucleotide compositions. Both mutation and selection biases, variously induced by replication, transcription and translation constraints, have been suggested to play a role in *B. burgdorferi* [16, 17] and more generally, across all prokaryotes. The loss of some DNA repair genes may also contribute to the low G+C content and heightened skew seen in *B. burgdorferi* [18, 19].

Thanks to the much expanded availability of complete genome sequences of bacterial species, it is now possible to perform large-scale comparative genomics studies [2, 4, 20]. A much larger number of bacterial genomes are in draft form, assembled to different levels of contiguity (contigs, scaffolds) and tentatively annotated using automated pipelines. Most of the existing methods for analyzing compositional biases and skews rely on fully or mostly contiguous genomic sequence and on the availability of detailed annotation of genes; such methods are much less applicable to the study of incomplete draft genomes.

We present here three novel metrics for quantitative analysis of genome skews. Our metrics are robust to assembly status and work well on incomplete genomes with draft annotation. Using these metrics, we analyzed a large collection of bacterial genomes—both complete and draft. We identified several groups of species and genera that present as outliers for one or more of the novel metrics. These outlier species are frequently pathogenic and tend to have unusual lifestyles, like *B. burgdorferi*.

## Methods

### Genomes studied

We obtained from the National Center for Biotechnology Information (NCBI) the genome sequence (in FASTA format) and current annotation (in General Feature Format, GFF) for 7948 bacterial species. We downloaded the “assembly_summary.txt” file from NCBI’s genome FTP site. This file provided various details on 86,822 genome assemblies including the organism name, RefSeq category (whether the genome considered “reference” for the species, “representative”, or otherwise) and assembly level (whether the genome is considered “completed”, or whether it is “incomplete”-assembled to chromosome, scaffold or contig level). Studying this file, we selected and downloaded:

1. 1581 “completed” genomes, (125 “reference”, 1456 “representative”),
2. 3303 “incomplete” genomes, (2 “reference”, 3301 “representative”), and
3. 3064 additional genomes, not repeating species names from the previous two sets, and prioritizing more advanced levels of completion where multiple assemblies are available for a given species.

For each genome, we included in the analysis all chromosomes, plasmids and sequence contigs at least 100 kb long. We removed from further analysis 210 genome assemblies for which the longest available sequence was shorter than 100 kb. The final set of genomes analyzed included 7738 assemblies.

#### Identification of origins of replication and terminator sites

For each sequence (chromosome, plasmid, scaffold and contig) in each genome assembly, we identified likely origins of replication and replication terminator sites using the GC disparity method [21], namely by identifying the minimum and maximum difference between the cumulative count of G and C along the genome. This method is independent of gene annotation and arbitrary window sizes; it can also efficiently determine the likely direction of replication for sequence fragments (scaffolds and contigs), whether or not they include an origin of replication or a terminator site. When the resulting origin or terminator site lay within 1% of either end of the sequence, we corrected the location to coincide with the nearest sequence end.

#### Segmentation and analysis

We used available gene annotation (in the GFF files) to segment each sequence 100 kb or longer into a series of contiguous and disjoint segments (of variable lengths) which can be genes (including CDS, tRNA, and rRNA) or intergenic segments. We stratified intergenic segments by considering the relative orientations of the flanking genes: between two genes in the same orientation, or between two genes in opposite orientations (“head to head” or “tail to tail”). Infrequently, consecutive gene segments may be annotated as overlapping. We treated the overlapping segments as intergenic.

We computed for each segment (genic or intergenic) its length, G+C content, GC skew, and TA skew. We further determined for each oriented segment (namely genes and intergenic segments between genes transcribed in the same orientation) whether their orientation is the same or opposite to the direction of replication, i.e., whether they are on the leading or lagging strand, relative to origin and terminator sites predicted as described above.

#### Computation of characteristic skews

Given a set of comparable segments in a genome assembly (e.g., all genes on the leading strand), we computed skews (GC skew = (G-C)/(G+C) and TA skew = (T-A)/(T+A)) for the set as the average of the corresponding individual segment skews, weighted by segment length. We thus compute four characteristic skews for each species: lead_GC_ and lead_TA_ for leading strand genes, and lag_GC_ and lag_TA_ for lagging strand genes. We also evaluated weighted medians instead of weighted averages, which yielded very similar results (not shown).

#### Computation of the cross-skew and dot-skew

The four characteristic skews for a species can be interpreted as two characteristic skew vectors: one for the leading strand genes (lead_GC_, lead_TA_) and the other for the lagging strand genes (lag_GC_, lag_TA_). We computed the cross-skew as:

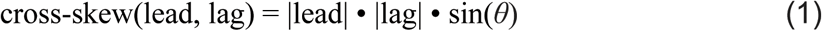

where |lead| = sqrt(lead_GC_^2^ + lead_TA_^2^), |lag| = sqrt(lag_GC_^2^ + lag_TA_^2^), and *θ* is the angle between the two vectors. Similarly, we computed the dot-skew as:

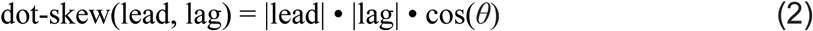

#### Computation of the residual skew

We modeled each of the four characteristic skews (lead_GC_, lead_TA_, lag_GC_ and lag_TA_) as a function of the G+C content for 7738 bacterial genome assemblies. For each characteristic skew we separated the genome assemblies with G+C content below or above 50% G+C (3635 and 4103 genomes, respectively), and fitted a robust regression line to each subset using the R function MASS∷lqs() (least trimmed sum of squares, [22]. We then computed a single skew deviation magnitude metric (the residual skew) for each genome as the root mean square deviation (RMSD) from the regression line across the four characteristic skews.

## Results

### The genomic skews of *B. burgdorferi* are anti-correlated

A map of the GC and TA skews in 1 kb bins along the main chromosome of *B. burgdorferi* B31 shows that most bins have either a strong GC skew or a strong TA skew (Fig. 1A). Most genomic bins show a combination of both types of skew, such that the sum of the two skews (G+T vs. C+A) appears to be almost constant; this is particularly evident when plotting the TA skew vs. the GC skew (Fig. 1B). Indeed, the strength of the two skews is complementary and negatively correlated. When expressing the skews relative to the leading strand (i.e., changing the sign of skews where the leading strand corresponds to the bottom strand), we observe that the combined skew has a narrower distribution (0.270 +/- 0.099) than expected from those of the individual skews (GC skew: 0.179 +/- 0.132; TA skew: 0.091 +/- 0.112; expected deviation for their combination: 0.173). The correlation between GC and TA skews is −0.68. The plasmids of *B. burgdorferi* also have significant skews [23].

**Figure 1.**
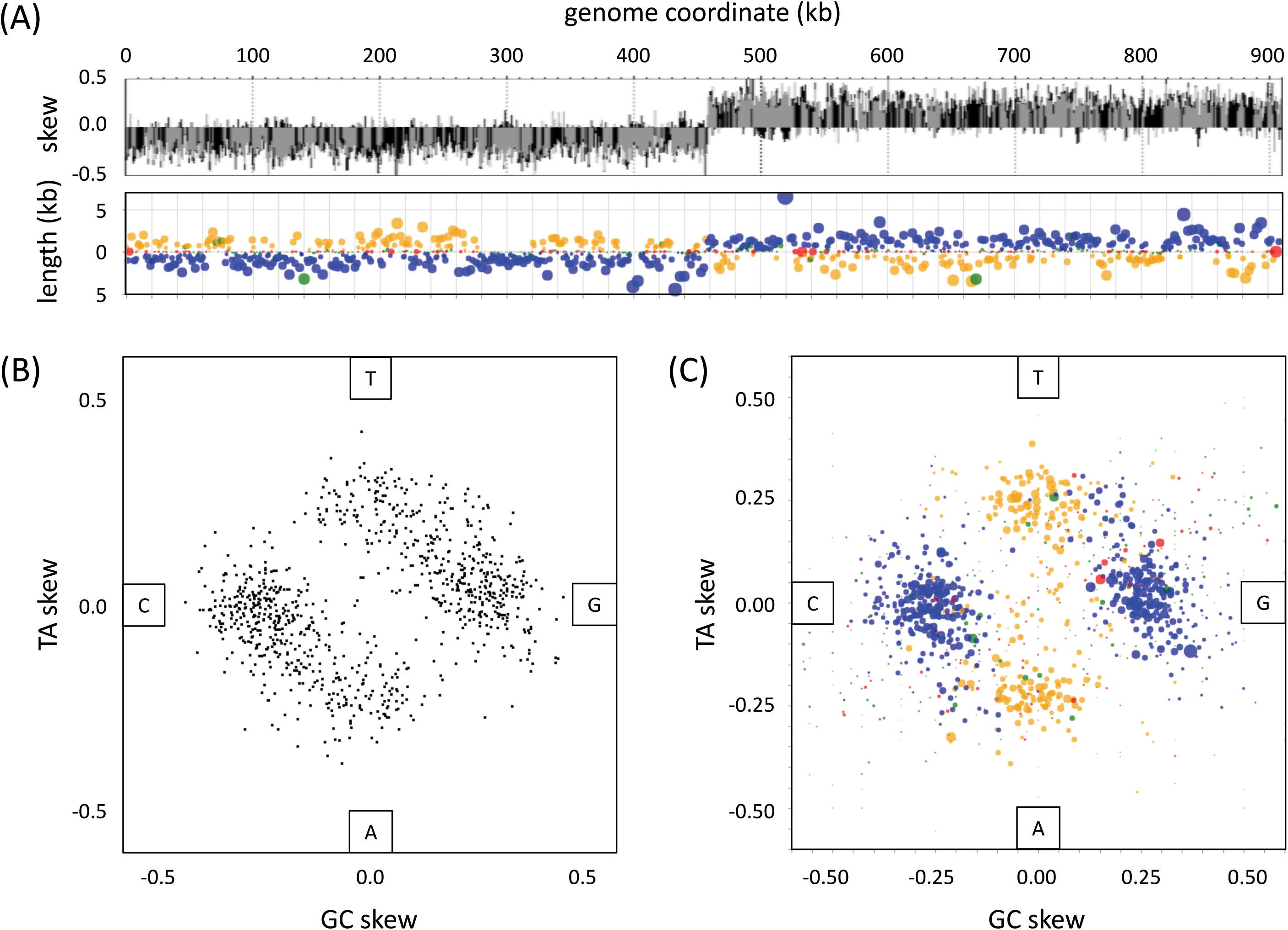
Bin-wise and gene-wise representations of genome skews in *B. burgdorferi.* A: TA (grey) and GC (black) skews in 1 kb bins along the (linear) main chromosome (top) and map of annotated genes and intergenic segments (bottom) showing strandedness and length of each segment: leading strand genes in blue, lagging strand genes in orange, intergenic segments flanked by genes in equal orientation in green, and intergenic segments flanked by genes in opposite orientations in red; circle area is proportional to segment length. The location of the origin of replication is evident from the sharp switch in skew sign from negative to positive. B: Comparison of skews per 1 kb bin. C: Comparison of skews per gene and intergenic segment; circle area is proportional to segment length.

In *B. burgdorferi*, the majority of genes are transcribed in the same direction as they are replicated (‘leading strand genes’, blue in Fig. 1) while some are transcribed in the direction opposite to replication (‘lagging strand genes’, orange in Fig. 1). Leading strand genes tend to display stronger GC skew (Fig. 1C), while lagging strand genes have strong TA skews. In intergenic segments (red and green in Fig. 1C), the two skews tend to be positively correlated.

#### The characteristic skews of *B. burgdorferi* genes

Since the clear symmetry in the skew comparison plot for *B. burgdorferi* (Fig. 1) reflects the opposite characteristics of the two halves of the chromosome (and likewise for each plasmid), each leading from the origin of replication to a terminator (or telomere, for a linear chromosome or plasmid), it is appropriate and convenient to express the skews relative to the leading strand orientation. This transformation simplifies the representation, showing two main clusters of genes corresponding to genes transcribed on the leading strand vs. on the lagging strand (Fig. 2). To further simplify the representation (aiming to enable comparison of a large set of genomes), these two clusters can be summarized by their average TA and GC skews, weighted by gene length (see Methods). We thus computed the four characteristic skews for *B. burgdorferi*: lead_GC_ = 0.259, lead_TA_ = 0.022, lag_GC_ = 0.016 and lag_TA_ = 0.211.

**Figure 2.**
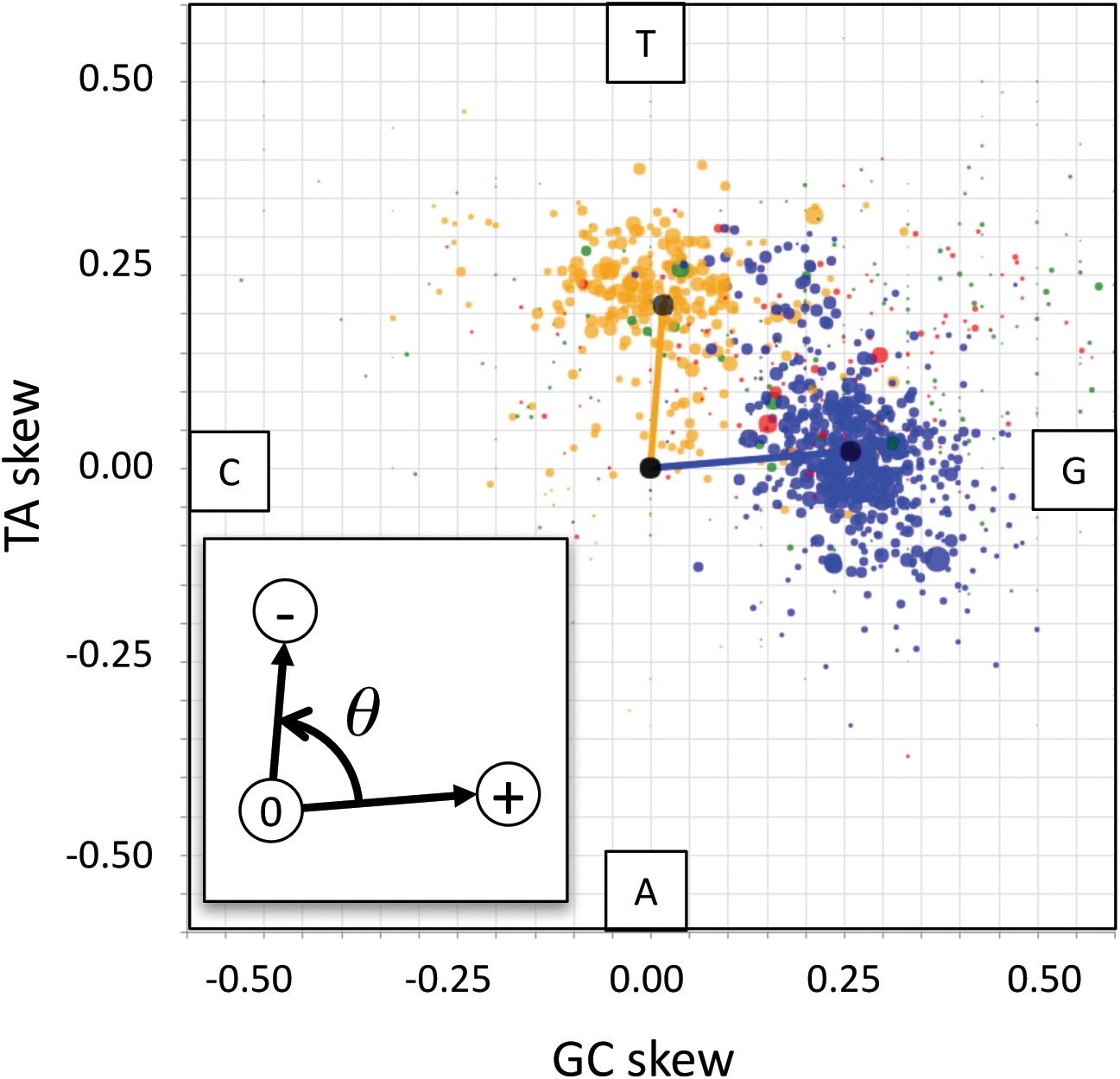
TA vs. GC skews of gene and intergenic segments oriented relative to the leading strand. Graphic elements as in Fig. 1C. The vectors point from the origin (zero skews) to the weighted average of skews for genes on the leading strand (+) and genes on the lagging strand (-). Inset: definition of the angle *θ* between the two vectors.

We visualized these skews as two vectors (leading, lagging) and computed the angle between them *θ* = 80.89° (Fig. 2, inset). Then, based on the length and angles of these two vectors (see Methods), we computed the *cross-skew* and *dot-skew* for *B. burgdorferi*: cross-skew(lead, lag) = 0.0584, dot-skew(lead, lag) = 0.0075. For other species within the Borrelia and Borreliella genera, these respectively ranged from 0.0562 to 0.0781 and from 0.0031 to 0.0293.

#### Learning from thousands of genomes

We similarly computed characteristic skews, angles and skew metrics for 7738 bacterial genome assemblies (see Methods). Visualization of these species-specific parameters demonstrates the wide diversity of bacterial genome composition; a few select examples are shown in Fig. 3. We observed genomes with strong skews and with negligible skews, at all possible angles between the skew vectors. We also created a web interface for generating species-specific skew plots and exploring their skew metrics, available at [24].

**Figure 3.**
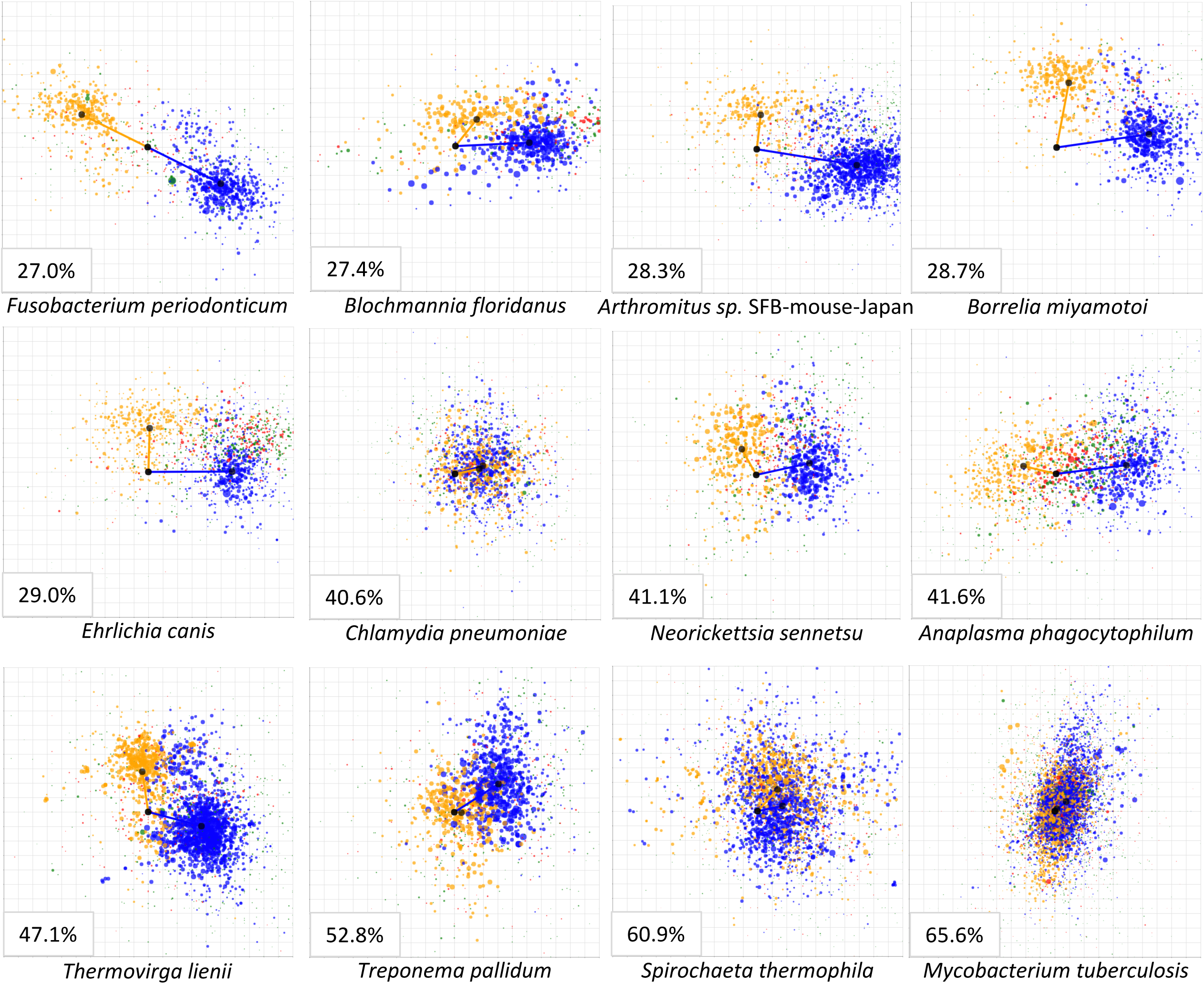
Examples of TA vs. GC skew plots for several bacterial species. Graphic elements as in Figs. 1C and 2. Each plot displays skews in the range [-0.5, 0.5]. Lower-left inset for each plot: average genomic G+C content for that species.

We compared the four characteristic skews of 7738 bacterial genome assemblies with their corresponding G+C content (Fig. 4). We observed that all skews are correlated with G+C content, and largely decrease in absolute value with increasing G+C content. These relationships are different for bacterial genomes with low vs. high G+C content. In fact, we observed a largely bimodal distribution of G+C content among sequenced bacterial genomes (Fig. 5, lower panel). We therefore fitted lines to the characteristic skews separately for bacterial genomes below and above 50% G+C content, and computed the deviations from the expected skews for each bacterial genome assembly. We used the R function MASS:lqs(), the leading robust linear regression method, to ensure that these lines accurately reflected the typical pattern, ignoring outliers.

**Figure 4.**
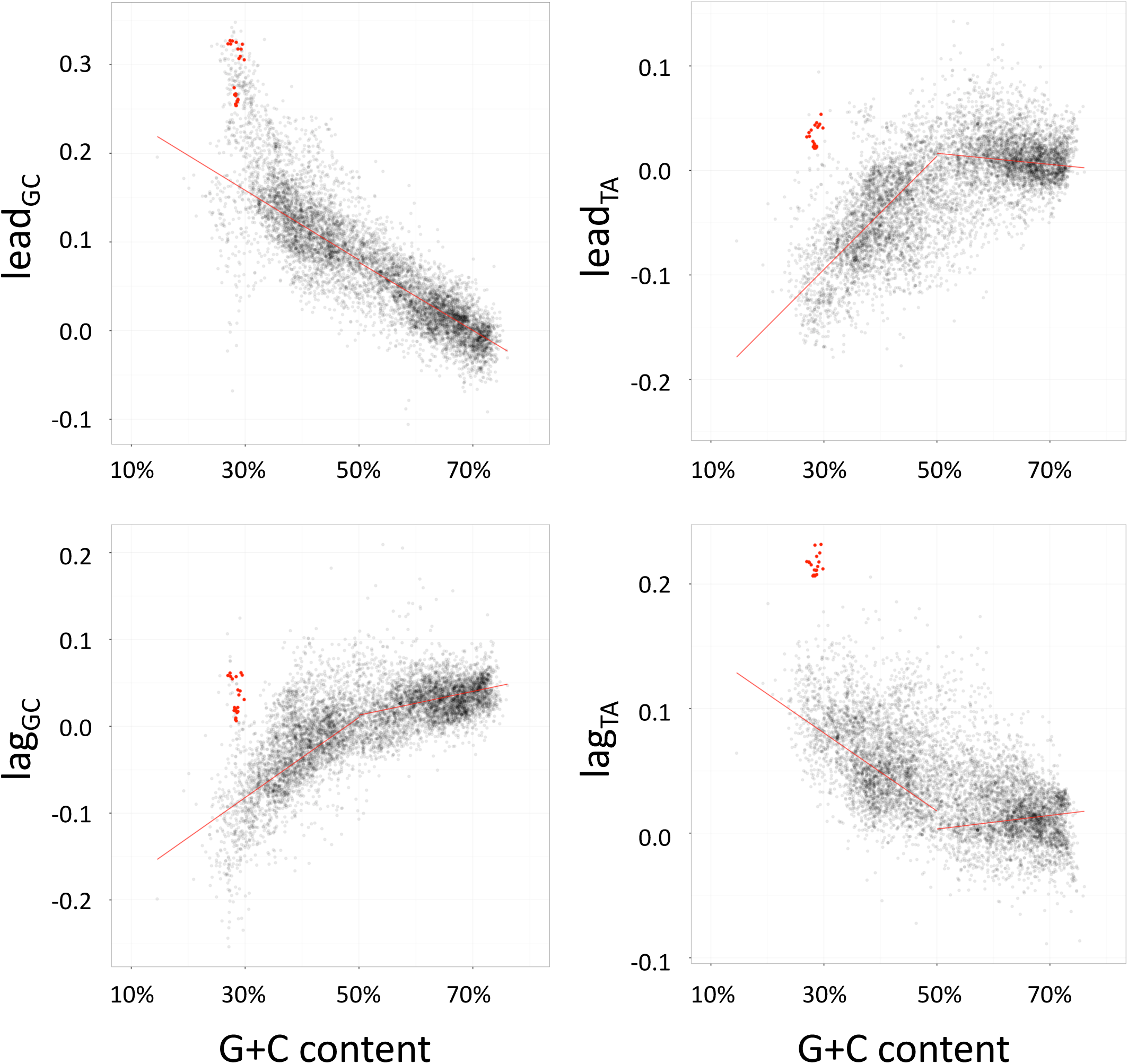
Relationship between the four characteristic skew values and G+C content, for 7738 bacterial genomes, highlighting Borreliaceae species (red points). Red lines represent robust regression lines computed by least quantile of squares method.

**Figure 5.**
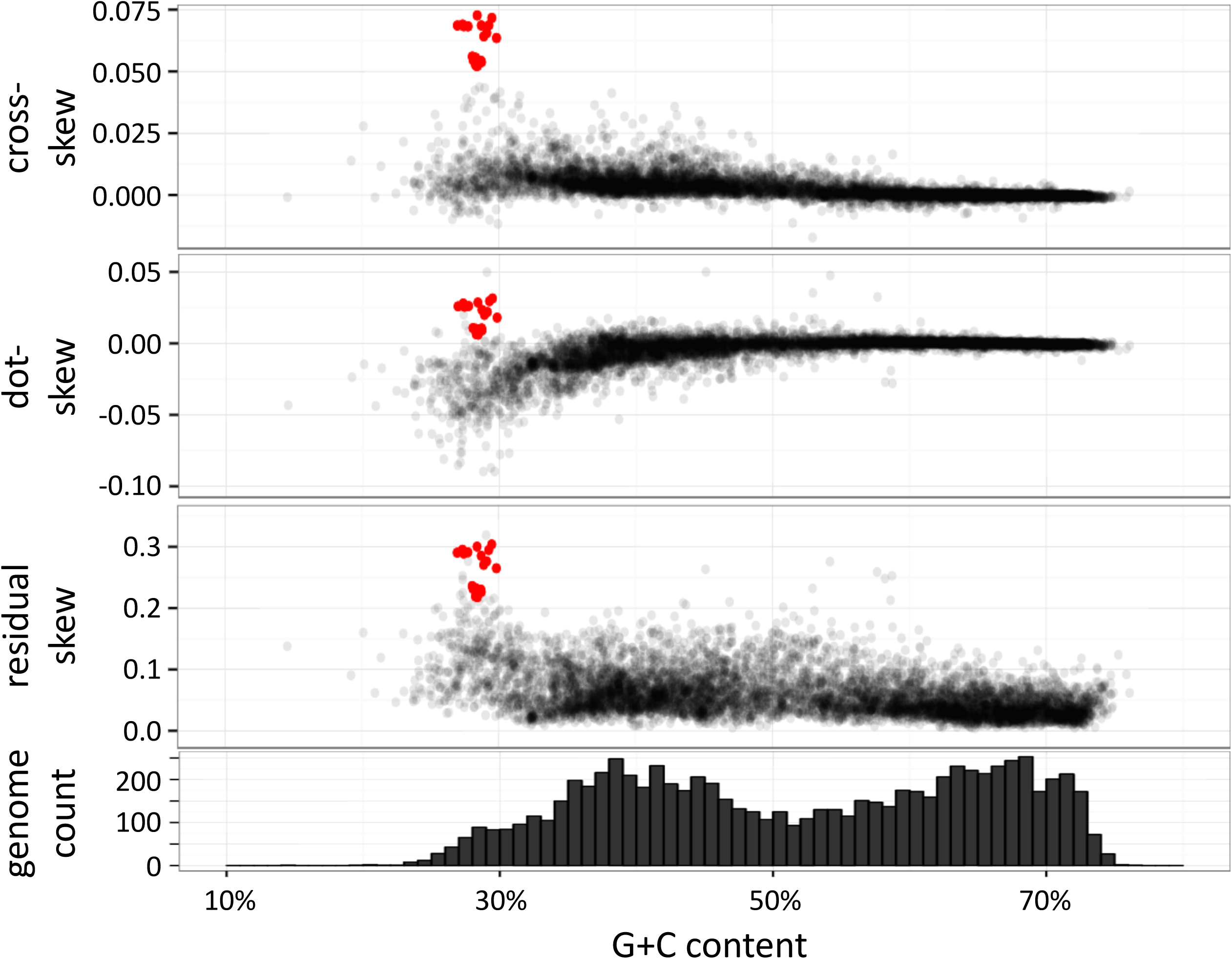
A: Skew metrics vs. G+C content for 7738 bacterial genomes, highlighting Borreliaceae species (red points). From top to bottom: cross-skew, dot-skew, residual skew, and histogram of number of species studied.

The lead_GC_ and lag_TA_ values of Borreliaceae genomes are large and are clear outliers relative to the entire data set of 7738 genomes (Fig. 4). On the other hand, while the Borreliaceae lead_TA_ and lag_GC_ are close to zero and are not outliers relative to the entire data set, they are unusual for bacterial species with low G+C content, which tend to have negative values for these characteristic skews (Fig. 4). Thus, Borreliaceae genomes are unusual for all four characteristic skews. The deviations of characteristic skews for *B. burgdorferi* from the skews predicted by the fitted lines at the G+C content for *B. burgdorferi* are 0.091, 0.120, 0.106 and 0.124 for lead_GC_, lead_TA_, lag_GC_ and lag_TA_, respectively. Borrelia species that cause relapsing fever have even larger deviations from the expected values.

#### Three novel metrics for analyzing genome skews

We described above several parameters for quantifying skews in individual bacterial genomes: the four characteristic skews and the magnitudes and angles of the vectors they define. Using these parameters, we defined two interrelated metrics for comparing and contrasting the skews of leading strand vs. lagging strand genes: the cross-skew and the dot-skew (see Methods). Furthermore, taking advantage of the availability of many thousand bacterial genome assemblies, we estimated expected values for each characteristic skew, as a function of the G+C content. We then used the observed deviations from these expected values to define a third metric: the *residual skew*.

We computed these three metrics for 7738 bacterial genome assemblies (available at [24]) and evaluated their relationship with G+C content (Fig. 5). For high G+C content bacteria, we observed that the cross-skew and the dot-skew are much more constrained than for lower G+C content species; these two metrics are most diverse for bacterial genomes under ~35% G+C. Compared to these two metrics, the residual skew is more diverse for all levels of G+C content. Borreliaceae genomes are clear outliers for all three metrics.

Finally, we combined all three metrics to generate a map of genome skews for all bacterial genomes (Fig. 6). In this map, most high G+C content bacteria are restricted to near the origin, while low G+C content bacteria show a more diverse spread. Borreliaceae genomes are seen as clear outliers, with the most extreme skew values corresponding to the group of Borrelia genomes that cause relapsing fever. Genomes in the genus Ehrlichia (see example in Fig. 3) are also outliers in all three metrics and show similar skew values as Borreliella genomes. Ehrlichia are intracellular vector-borne pathogens of vertebrates; like Borrelia, they have diminished biosynthetic abilities [25]. Ehrlichia are in the Rickettsiales order and are phylogenetically unrelated to Borreliaceae; the genome of *Ehrlichia canis* has a single circular chromosome and no plasmids [26]. Multiple other genera became evident as outliers of interest, discussed below.

**Figure 6.**
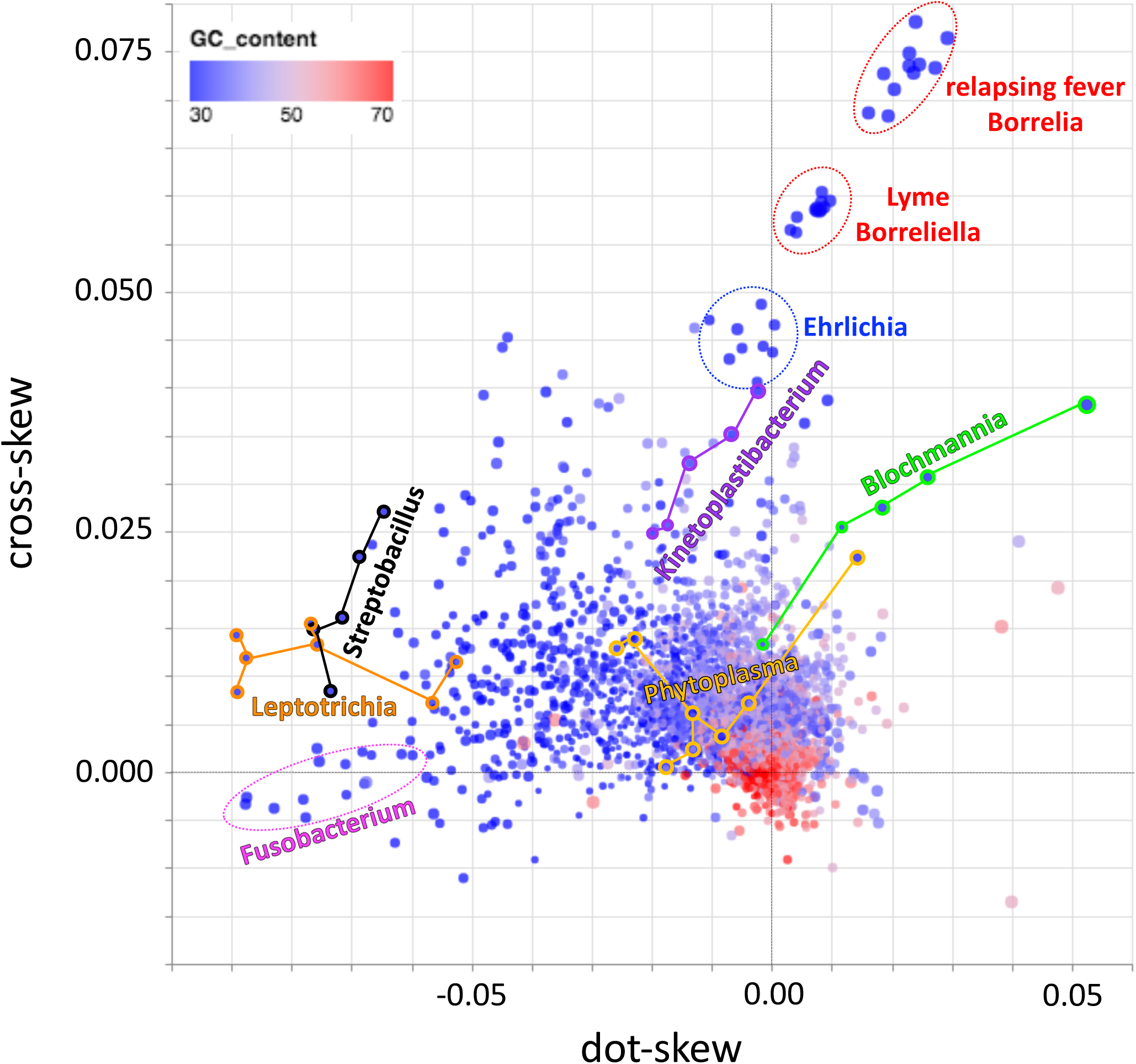
Integration of skew metrics (cross-skew vs. dot-skew, point size represents residual skew) for 7738 bacterial genomes, highlighting some genera of interest. All genomes colored by G+C content, ranging from low (blue) to high (red).

## Discussion

We have devised three novel metrics to study bacterial genome composition biases, integrating knowledge of the nucleotide skews in annotated genes, the direction of transcription relative to replication, and the G+C content of the genome.

The first two metrics (cross-skew and dot-skew) are computed based on knowledge of an individual genome’s characteristic skew vectors, and they quantify the strength and relationship between the mutation and selection pressures on genes on the leading vs. lagging strands.

Positive dot-skew values (Fig. 6, top) indicate similar compositional constraints on all genes, relative to the direction of replication; an example of this pattern is observed in the obligate intracellular parasite *Chlamydia pneumoniae* [27] (Fig. 3). Conversely, negative dot-skew values (Fig. 6, bottom) reflect opposite compositional constraints on leading and lagging strand genes (i.e., transcribed in the same or opposite direction as they are replicated); extreme examples of this pattern are observed in fusobacteria including *Fusobacterium periodonticum* [28], *Leptotrichia buccalis [29]*, and *Streptobacillus moniliformis [30]*, the causal agent of rat bite fever. Positive dot-skew values can thus be interpreted as reflecting constraints driven mostly by the replication process, while negative dot-skew values largely reflect transcriptional and translational constraints [4].

The cross-skew quantifies the strength and orthogonality of the compositional skew vectors for leading and lagging strand genes. Genomes with high cross-skew values (Fig. 6, right) demonstrate skew patterns inconsistent with purely replicational or transcriptional constraints; Borreliaceae and Ehrlichia species are prime examples of this pattern.

Borreliaceae and Ehrlichia species lack amino acid and nucleotide synthesis pathways; the observed skew patterns in these pathogens may thus reflect a relaxation of the selection for energy efficiency that drives nucleotide usage and thus skews [4], possibly combined with more complex constraints imposed by the a life cycle that involves recurring transitions between mammalian and invertebrate (tick) hosts. We observed similar skew patterns in Kinetoplastibacteria (Fig. 6), which are endosymbionts of insect-infecting trypanosomatid flagellates [31] with multiple biosynthetic adaptations to life in the intracellular environment. Likewise, we observed distinct skew patterns among Blochmannia species (Fig. 6); these are also intracellular endosymbionts that lost multiple biosynthetic pathways and rely on the metabolic machinery of their carpenter ant hosts [32].

The third metric (residual skew) capitalizes on the current availability of thousands of complete or draft bacterial genomes to empirically assess how unusual a genome’s skews are relative to the expected values as learned from other genomes. This analysis, which has not been possible until recent times, revealed that bacterial genomes with low G+C content typically have negative TA skews in leading strand genes and negative GC skews in lagging strand genes, and that these negative skews increase in magnitude as G+C content decreases (Fig. 4). On the background of these trends, the weakly positive skews observed in Borreliaceae species are highly unusual. This pattern is not evident relative to the global collection of genomes since the weakly positive Borreliaceae skews are comparable to those observed in high G+C content bacteria. Our regression analysis quantifies these deviations from expectation and integrates them into a unified metric that highlights the unusual skews in Borreliaceae species (Fig. 5) and also identifies other species as having skew patterns that are significantly unusual relative to the bulk of bacterial species. Of particular note are Phytoplasma species (Fig. 6); these are intracellular pathogens of multiple plant species that use insects as transmission vectors [33, 34], in similarity to Borreliaceae and Ehrlichia for mammals.

## Conclusions

We described here three novel metrics for quantifying bacterial genome composition skews and presented examples of their application to identify bacterial species with unusual skew patterns. Our metrics take advantage both of information about the genome of a single species and of patterns discernable from studying genomes of thousands of species - even those not yet finished and fully annotated. While some of the genera identified as skew outliers are phylogenetically close (e.g., Fusobacterium, Streptobacillus and Leptotrichia), our metrics identified similar skew patterns in genera of bacteria that are phylogenetically unrelated, like Borrelia, Ehrlichia and Kinetoplastibacterium, and (when considering the residual skew) Phytoplasma. These very disparate bacterial species share lifestyle characteristics (intracellularity and transmission via insect vectors), suggesting that our novel metrics successfully capture effects on genome composition of biosynthetic constraints and of interaction with the hosts.

## List of abbreviations

GFF: General Feature Format
NCBI: National Center for Biotechnology Information
RMSD: root mean square deviation

## Declarations

### Ethics approval and consent to participate

Not applicable.

### Consent for publication

Not applicable.

### Availability of data and material

Software for computing skews for individual genomes, regression RMSD parameters, a table of skew characteristics for the 7738 bacterial genomes, and Vega-Lite [35] interactive visualizations are available at [24]. The table of characteristics for 7738 bacterial genomes is also included as supplementary material.

### Authors’ contributions

GG conceived of the study. LMJC, MR and GG designed the methods and implemented the software. LMJC, NJ and GG performed analyses. LMJC, CL, MR and GG wrote the manuscript. All authors approved its final version.

## Acknowledgements

We wish to thank Arian Smit and Jeff Boore for helpful discussions.

